## Supplementary Material for "A novel multiplex RNAi therapy simultaneously targets *Hif1a* and *Hif2a* to defy retinal degeneration in two models of AMD"

Figure S1

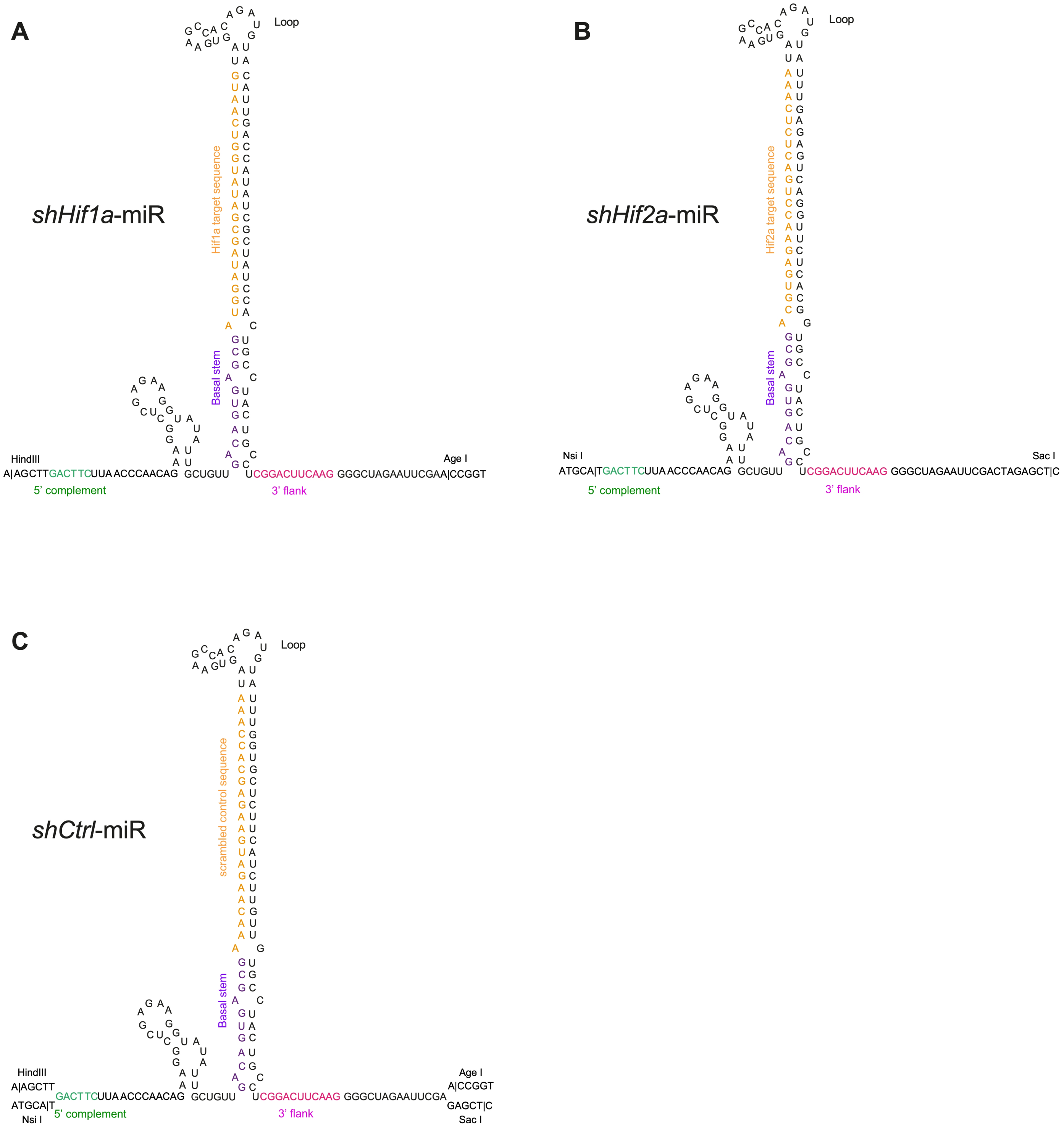

A
shHif1a-miR
Loop
Hif1a target sequence
Basal stem
HindIII
Age I
A|AGCTTGACTTCUUA ACCCAACAG GCUGUU UCGGACUUCAAG GGGCUAGAAUUCGAA|CCGGT
5' complement
3' flank
B
shHif2a-miR
Loop
Hif2a target sequence
Basal stem
Nsi I
Sac I
ATGCA|TGACTTCUUA ACCCAACAG GCUGUU UCGGACUUCAAG GGGCUAGAAUUCGACTAGAGCT|C
5' complement
3' flank
C
shCtrl-miR
Loop
scrambled control sequence
Basal stem
HindIII
A|AGCTT
ATGCA|T
Nsi I
GACTTCUUA ACCCAACAG GCUGUU UCGGACUUCAAG GGGCUAGAAUUCGA
Age I
A|CCGGT
GAGCT|C
Sac I
5' complement
3' flank

Figure S2

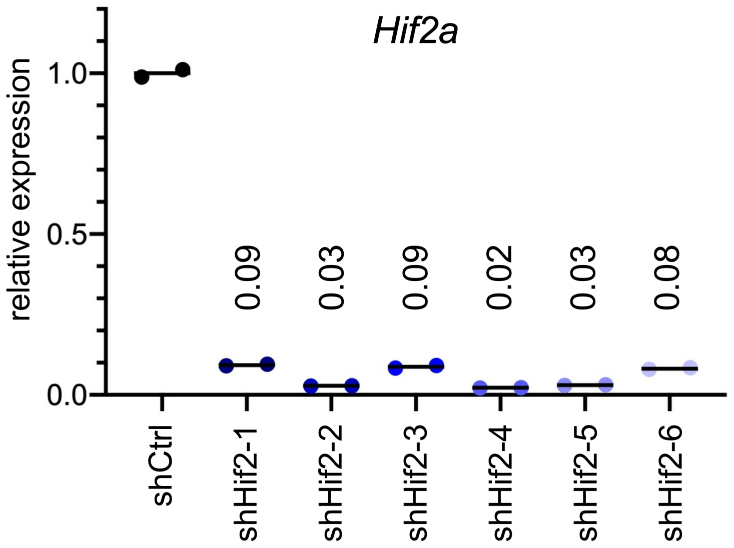
Hif2a
relative expression
1.0
0.5
0.0
0.09
0.03
0.09
0.02
0.03
0.08
shCtrl
shHif2-1
shHif2-2
shHif2-3
shHif2-4
shHif2-5
shHif2-6

Figure S3

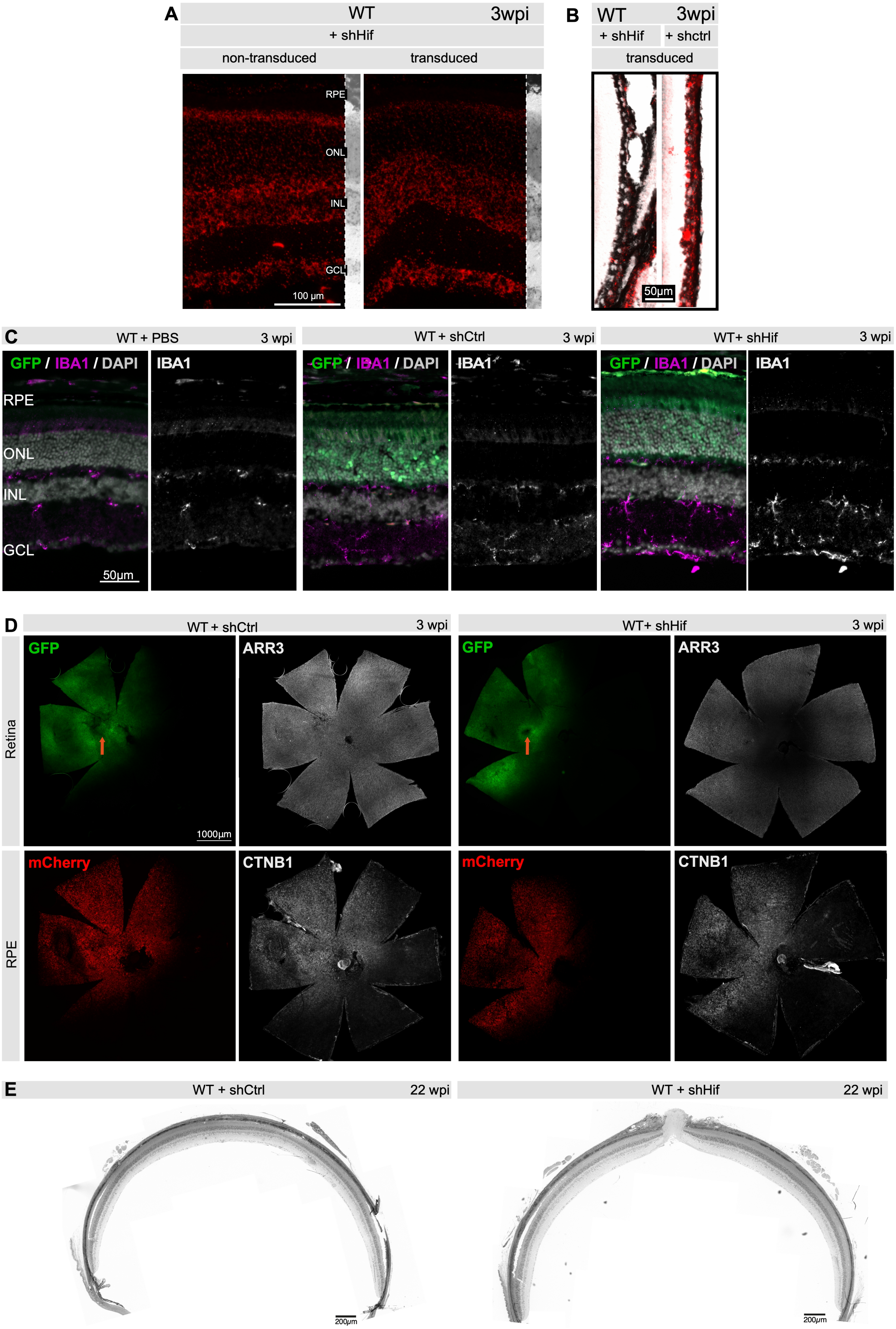
A
WT
3wpi
+ shHif
non-transduced
transduced
RPE
ONL
INL
GCL
100 µm
B
WT
3wpi
+ shHif
+ shctrl
transduced
50µm
C
WT + PBS
3 wpi
WT + shCtrl
3 wpi
WT+ shHif
3 wpi
GFP / IBA1 / DAPI
IBA1
RPE
ONL
INL
GCL
50µm
D
WT + shCtrl
3 wpi
WT+ shHif
3 wpi
Retina
RPE
GFP
ARR3
1000µm
mCherry
CTNB1
E
WT + shCtrl
22 wpi
WT + shHif
22 wpi
200µm

Figure S4

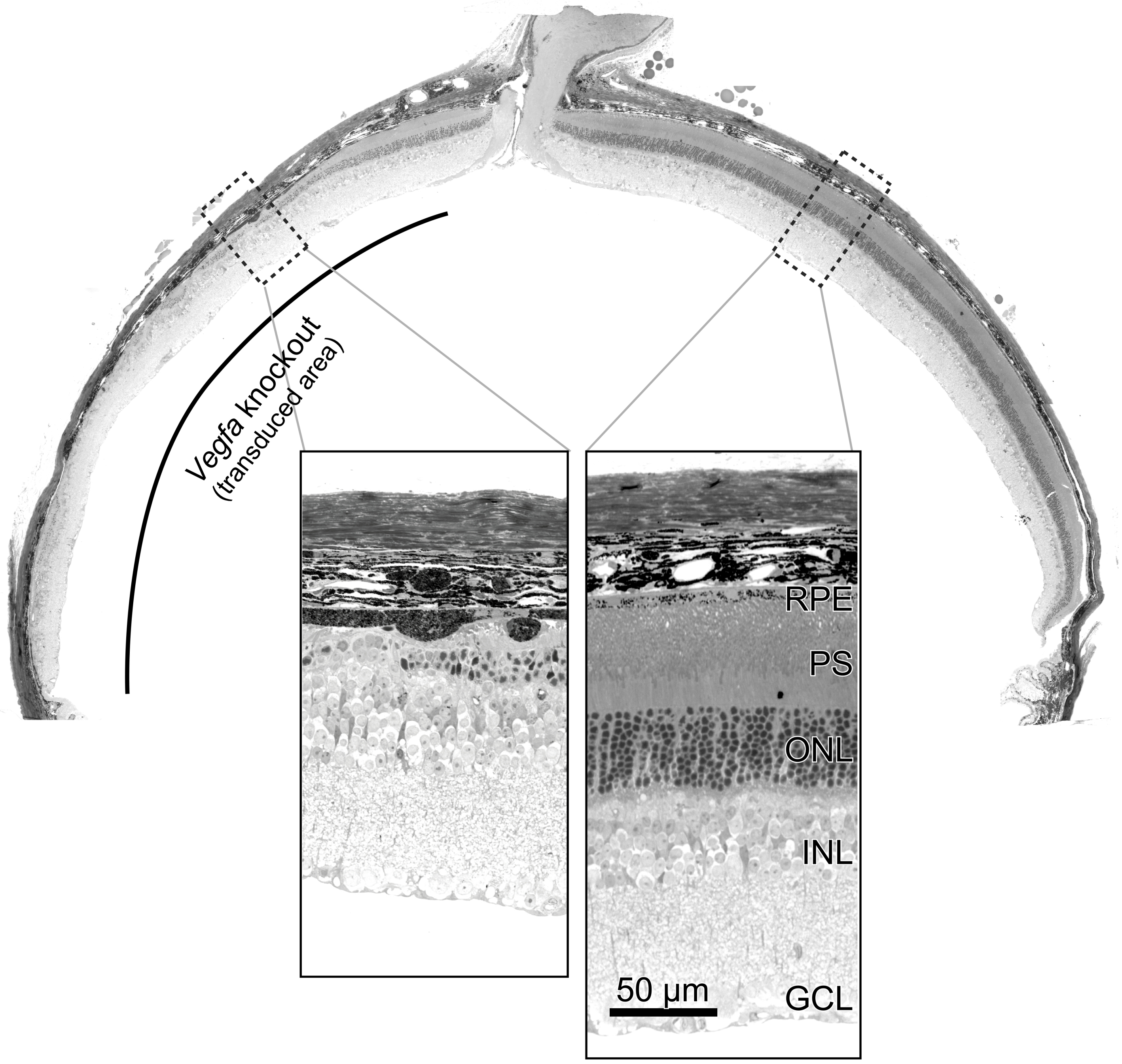

### Supplemental data file SD1

#### Plasmid sequence *shHif*

GGTTGGCCACTCCCTCTCTGCGCGCTCGCTCGCTCACTGAGGCCGGGCGACCAAAGGTCGCCCGACGCCCGGGCTTTGCC
CGGGCGGCCTCAGTGAGCGAGCGAGCGCGCAGAGAGGGAGTGGCCAACTCCATCACTAGGGGTTCCTAGATCTGAATTCG
GTACCGGGCCCCAGAAGCCTGGTGGTTGTTTGTCCTTCTCAGGGGAAAAGTGAGGCGGCCCCTTGGAGGAAGGGGCCGGG
CAGAATGATCTAATCGGATTCCAAGCAGCTCAGGGGATTGTCTTTTTCTAGCACCTTCTTGCCACTCCTAAGCGTCCTCC
GTGACCCCGGCTGGGATTTAGCCTGGTGCTGTGTCAGCCCCGGTCTCCCAGGGGCTTCCCAGTGGTCCCCAGGAACCCTC
GACAGGGCCCGGTCTCTCTCGTCCAGCAAGGGCAGGGACGGGCCACAGGCCAAGGGCTCTAGAGGATCCGGTACTCGAGG
AACTGAAAAACCAGAAAGTTAACTGGTAAGTTTAGTCTTTTTGTCTTTTATTTCAGGTCCCGGATCCGGTGGTGGTGCAA
ATCAAAGAACTGCTCCTCAGTGGATGTTGCCTTTACTTCTAGGCCTGTACGGAAGTGTTACTTCTGCTCTAAAAGCTGCG
GAATTGTACCCGCGGCCGCCGCCACCATGAGCAAGGGCGAGGAACTGTTCACTGGCGTGGTCCCAATTCTCGTGGAACTG
GATGGCGATGTGAATGGGCACAAATTTTCTGTCAGCGGAGAGGGTGAAGGTGATGCCACATACGGAAAGCTCACCCTGAA
ATTCATCTGCACCACTGGAAAGCTCCCTGTGCCATGGCCAACACTGGTCACTACCCTGACCTATGGCGTGCAGTGCTTTT
CCAGATACCCAGACCATATGAAGCAGCATGACTTTTTCAAGAGCGCCATGCCCGAGGGCTATGTGCAGGAGAGAACCATC
TTTTTCAAAGATGACGGGAACTACAAGACCCGCGCTGAAGTCAAGTTCGAAGGTGACACCCTGGTGAATAGAATCGAGCT
GAAGGGCATTGACTTTAAGGAGGATGGAAACATTCTCGGCCACAAGCTGGAATACAACTATAACTCCCACAATGTGTACA
TCATGGCCGACAAGCAAAAGAATGGCATCAAGGTCAACTTCAAGATCAGACACAACATTGAGGATGGATCCGTGCAGCTG
GCCGACCATTATCAACAGAACACTCCAATCGGCGACGGCCCTGTGCTCCTCCCAGACAACCATTACCTGTCCACCCAGTC
TGCCCTGTCTAAAGATCCCAACGAAAAGAGAGACCACATGGTCCTGCTGGAGTTTGTGACCGCTGCTGGGATCACACATG
GCATGGACGAGCTGTACAAGTGAGCGGCCGCAAGCTTGACTTCTTAACCCAACAGAAGGCTCGAGAAGGTATATTGCTGT
TGACAGTGAGCG**ATGGATAGCGATATGGTCAATG**TAGTGAAGCCACAGATGTACATTGACCATATCGCTATCCACTGCCT
ACTGCCTCGGACTTCAAGGGGCTAGAATTCGAACCGGTTTCCTAGGAGACATGATAAGATACATTGATGAGTTTGGACAA
ACCACAACTAGAATGCAGTGAAAAAAATGCTTTATTTGTGAAATTTGTGATGCTATTGCTTTATTTGTAACCATTATAAG
CTGCAATAAACAAGTTAACAACAACAATTGCATTCATTTTATGTTTCAGGTTCAGGGGGAGGTGTGGGAGGTTTTTTACG
CGTCAATTCTGTCATTTTACTAGGGTGATGAAATTCCCAAGCAACACCATCCTTTTCAGATAAGGGCACTGAGGCTGAGA
GAGGAGCTGAAACCTACCCGGCGTCACCACACACAGGTGGCAAGGCTGGGACCAGAAACCAGGACTGTTGACTGCAGCCC
GGTATTCATTCTTTCCATAGCCCACAGGGCTGTCAAAGACCCCAGGGCCTAGTCAGAGGCTCCTCCTTCCTGGAGAGTTC
CTGGCACAGAAGTTGAAGCTCAGCACAGCCCCCTAACCCCCAACTCTCTCTGCAAGGCCTCAGGGGTCAGAACACTGGTG
GAGCAGATCCTTTAGCCTCTGGATTTTAGGGCCATGGTAGAGGGGGTGTTGCCCTAAATTCCAGCCCTGGTCTCAGCCCA
ACACCCTCCAAGAAGAAATTAGAGGGGCCATGGCCAGGCTGTGCTAGCCGTTGCTTCTGAGCAGATTACAAGAAGGGACC
AAGACAAGGACTCCTTTGTGGAGGTCCTGGCTTAGGGAGTCAAGTGACGGCGGCTCAGCACTCACGTGGGCAGTGCCAGC
CTCTAAGAGTGGGCAGGGGCACTGGCCACAGAGTCCCAGGGAGTCCCACCAGCCTAGTCGCCAGACCGGGGATCCTCTAG
AGGATCCGGTACTCGAGGAACTGAAAAACCAGAAAGTTAACTGGTAAGTTTAGTCTTTTTGTCTTTTATTTCAGGTCCCG
GATCCGGTGGTGGTGCAAATCAAAGAACTGCTCCTCAGTGGATGTTGCCTTTACTTCTAGGCCTGTACGGAAGTGTTACT
TCTGCTCTAAAAGCTGCGGAATTGTACCCGCGGCCGCCACCATGGTGAGCAAGGGCGAGGAGGACAACATGGCCATCATC
AAGGAGTTCATGCGCTTCAAGGTGCACATGGAGGGCTCCGTGAACGGCCACGAGTTCGAGATCGAGGGCGAGGGCGAGGG
CCGCCCCTACGAGGGCACCCAGACCGCCAAGCTGAAGGTGACCAAGGGCGGCCCCCTGCCCTTCGCCTGGGACATCCTGT
CCCCTCAGTTCATGTACGGCTCCAAGGCCTACGTGAAGCACCCCGCCGACATCCCCGACTACTTGAAGCTGTCCTTCCCC
GAGGGCTTCAAGTGGGAGCGCGTGATGAACTTCGAGGACGGCGGCGTGGTGACCGTGACCCAGGACTCCTCCCTGCAGGA
CGGCGAGTTCATCTACAAGGTGAAGCTGCGCGGCACCAACTTCCCCTCCGACGGCCCCGTAATGCAGAAGAAGACCATGG
GCTGGGAGGCCTCCTCCGAGCGGATGTACCCCGAGGACGGCGCCCTGAAGGGCGAGATCAAGCAGAGGCTGAAGCTGAAG
GACGGCGGCCACTACGACGCCGAGGTCAAGACCACCTACAAGGCCAAGAAGCCCGTGCAGCTGCCCGGCGCCTACAACGT
CAACATCAAGCTGGACATCACCTCCCACAACGAGGACTACACCATCGTGGAACAGTACGAGCGCGCCGAGGGCCGCCACT
CCACCGGCGGCATGGACGAGCTGTACAAGTAAATGCATGACTTCTTAACCCAACAGAAGGCTCGAGAAGGTATATTGCTG
TTGACAGTGAGCG**ACGTGAGAACCTGACTCTCAAA**TAGTGAAGCCACAGATGTATTTGAGAGTCAGGTTCTCACGGTGCC
TACTGCCTCGGACTTCAAGGGGCTAGAATTCGACTAGAGCTCGCTGATCAGCCTCGACTGTGCCTTCTAGTTGCCAGCCA
TCTGTTGTTTGCCCCTCCCCCGTGCCTTCCTTGACCCTGGAAGGTGCCACTCCCACTGTCCTTTCCTAATAAAATGAGGA
AATTGCATCGCATTGTCTGAGTAGGTGTCATTCTATTCTGGGGGGTGGGGTGGGGCAGGACAGCAAGGGGGAGGATTGGG
AAGACAATAGCAGGCATGCTGGGGAGAGATCTAGGAACCCCTAGTGATGGAGTTGGCCACTCCCTCTCTGCGCGCTCGCT
CGCTCACTGAGGCCGCCCGGGCAAAGCCCGGGCGTCGGGCGACCTTTGGTCGCCCGGCCTCAGTGAGCGAGCGAGCGCGC
AGAGAGGGAGTGGCCAACCCCCCCCCCCCCCCCCCCTGCAGCCCTGCATTAATGAATCGGCCAACGCGCGGGGAGAGGCGG
TTTGCGTATTGGGCGCTCTTCCGCTTCCTCGCTCACTGACTCGCTGCGCTCGGTCGTTCGGCTGCGGCGAGCGGTATCAG
CTCACTCAAAGGCGGTAATACGGTTATCCACAGAATCAGGGGATAACGCAGGAAAGAACATGTGAGCAAAAGGCCAGCAA
AAGGCCAGGAACCGTAAAAAGGCCGCGTTGCTGGCGTTTTTCCATAGGCTCCGCCCCCCTGACGAGCATCACAAAAATCG
ACGCTCAAGTCAGAGGTGGCGAAACCCGACAGGACTATAAAGATACCAGGCGTTTCCCCCTGGAAGCTCCCTCGTGCGCT
CTCCTGTTCCGACCCTGCCGCTTACCGGATACCTGTCCGCCTTTCTCCCTTCGGGAAGCGTGGCGCTTTCTCAATGCTCA
CGCTGTAGGTATCTCAGTTCGGTGTAGGTCGTTCGCTCCAAGCTGGGCTGTGTGCACGAACCCCCCGTTCAGCCCGACCG
CTGCGCCTTATCCGGTAACTATCGTCTTGAGTCCAACCCGGTAAGACACGACTTATCGCCACTGGCAGCAGCCACTGGTA
ACAGGATTAGCAGAGCGAGGTATGTAGGCGGTGCTACAGAGTTCTTGAAGTGGTGGCCTAACTACGGCTACACTAGAAGG
ACAGTATTTGGTATCTGCGCTCTGCTGAAGCCAGTTACCTTCGGAAAAAGAGTTGGTAGCTCTTGATCCGGCAAACAAAC
CACCGCTGGTAGCGGTGGTTTTTTTGTTTGCAAGCAGCAGATTACGCGCAGAAAAAAAGGATCTCAAGAAGATCCTTTGA
TCTTTTCTACGGGGTCTGACGCTCAGTGGAACGAAAACTCACGTTAAGGGATTTTGGTCATGAGATTATCAAAAAGGATC

TTCACCTAGATCCTTTTAAATTAAAAATGAAGTTTTAAATCAATCTAAAGTATATATGAGTAAACTTGGTCTGACAGTTA
CCAATGCTTAATCAGTGAGGCACCTATCTCAGCGATCTGTCTATTTCGTTCATCCATAGTTGCCTGACTCCCCGTCGTGT
AGATAACTACGATACGGGAGGGCTTACCATCTGGCCCCAGTGCTGCAATGATACCGCGAGACCCACGCTCACCGGCTCCA
GATTTATCAGCAATAAACCAGCCAGCCGGAAGGGCCGAGCGCAGAAGTGGTCCTGCAACTTTATCCGCCTCCATCCAGTC
TATTAATTGTTGCCGGGAAGCTAGAGTAAGTAGTTCGCCAGTTAATAGTTTGCGCAACGTTGTTGCCATTGCTACAGGCA
TCGTGGTGTCACGCTCGTCGTTTGGTATGGCTTCATTCAGCTCCGGTTCCCAACGATCAAGGCGAGTTACATGATCCCCC
ATGTTGTGCAAAAAAGCGGTTAGCTCCTTCGGTCCTCCGATCGTTGTCAGAAGTAAGTTGGCCGCAGTGTTATCACTCAT
GGTTATGGCAGCACTGCATAATTCTCTTACTGTCATGCCATCCGTAAGATGCTTTTCTGTGACTGGTGAGTACTCAACCA
AGTCATTCTGAGAATAGTGTATGCGGCGACCGAGTTGCTCTTGCCCGGCGTCAATACGGGATAATACCGCGCCACATAGC
AGAACTTTAAAAGTGCTCATCATTGGAAAACGTTCTTCGGGGCGAAAACTCTCAAGGATCTTACCGCTGTTGAGATCCAG
TTCGATGTAACCCACTCGTGCACCCAACTGATCTTCAGCATCTTTTACTTTCACCAGCGTTTCTGGGTGAGCAAAAACAG
GAAGGCAAAATGCCGCAAAAAAGGGAATAAGGGCGACACGGAAATGTTGAATACTCATACTCTTCCTTTTTCAATATTAT
TGAAGCATTTATCAGGGTTATTGTCTCATGAGCGGATACATATTTGAATGTATTTAGAAAAATAAACAAATAGGGGTTCC
GCGCACATTTCCCCGAAAAGTGCCACCTGACGTCTAAGAAACCATTATTATCATGACATTAACCTATAAAAATAGGCGTA
TCACGAGGCCCTTTCGTCTCGCGCGTTTCGGTGATGACGGTGAAAACCTCTGACACATGCAGCTCCCGGAGACGGTCACA
GCTTGTCTGTAAGCGGATGCCGGGAGCAGACAAGCCCGTCAGGGCGCGTCAGCGGGTGTTGGCGGGTGTCGGGGCTGGCT
TAACTATGCGGCATCAGAGCAGATTGTACTGAGAGTGCACCATATGCGGTGTGAAATACCGCACAGATGCGTAAGGAGAA
AATACCGCATCAGGAAATTGTAAACGTTAATATTTTGTTAAAATTCGCGTTAAATTTTTGTTAAATCAGCTCATTTTTTA
ACCAATAGGCCGAAATCGGCAAAATCCCTTATAAATCAAAAGAATAGACCGAGATAGGGTTGAGTGTTGTTCCAGTTTGG
AACAAGAGTCCACTATTAAAGAACGTGGACTCCAACGTCAAAGGGCGAAAAACCGTCTATCAGGGCGATGGCCCACTACG
TGAACCATCACCCTAATCAAGTTTTTTGGGGTCGAGGTGCCGTAAAGCACTAAATCGGAACCCTAAAGGGAGCCCCCGAT
TTAGAGCTTGACGGGGAAAGCCGGCGAACGTGGCGAGAAAGGAAGGGAAGAAAGCGAAAGGAGCGGGCGCTAGGGCGCTG
GCAAGTGTAGCGGTCACGCTGCGCGTAACCACCACACCCGCCGCGCTTAATGCGCCGCTACAGGGCGCGTCGCGCCATTC
GCCATTCAGGCTACGCAACTGTTGGGAAGGGCGATCGGTGCGGGCCTCTTCGCTATTACGCCAGGCTGCAGGGGGGGGGG
GGGGGG

**shHif1a:** **ATGGATAGCGATATGGTCAATG**

**shHif2a:** **ACGTGAGAACCTGACTCTCAAA**

#### Plasmid sequence *shCtrl*

GGTTGGCCACTCCCTCTCTGCGCGCTCGCTCGCTCACTGAGGCCGGGCGACCAAAGGTCGCCCGACGCCCGGGCTTTGCC
CGGGCGGCCTCAGTGAGCGAGCGAGCGCGCAGAGAGGGAGTGGCCAACTCCATCACTAGGGGTTCCTAGATCTGAATTCG
GTACCGGGCCCCAGAAGCCTGGTGGTTGTTTGTCCTTCTCAGGGGAAAAGTGAGGCGGCCCCTTGGAGGAAGGGGCCGGG
CAGAATGATCTAATCGGATTCCAAGCAGCTCAGGGGATTGTCTTTTTCTAGCACCTTCTTGCCACTCCTAAGCGTCCTCC
GTGACCCCGGCTGGGATTTAGCCTGGTGCTGTGTCAGCCCCGGTCTCCCAGGGGCTTCCCAGTGGTCCCCAGGAACCCTC
GACAGGGCCCGGTCTCTCTCGTCCAGCAAGGGCAGGGACGGGCCACAGGCCAAGGGCTCTAGAGGATCCGGTACTCGAGG
AACTGAAAAACCAGAAAGTTAACTGGTAAGTTTAGTCTTTTTGTCTTTTATTTCAGGTCCCGGATCCGGTGGTGGTGCAA
ATCAAAGAACTGCTCCTCAGTGGATGTTGCCTTTACTTCTAGGCCTGTACGGAAGTGTTACTTCTGCTCTAAAAGCTGCG
GAATTGTACCCGCGGCCGCCGCCACCATGAGCAAGGGCGAGGAACTGTTCACTGGCGTGGTCCCAATTCTCGTGGAACTG
GATGGCGATGTGAATGGGCACAAATTTTCTGTCAGCGGAGAGGGTGAAGGTGATGCCACATACGGAAAGCTCACCCTGAA
ATTCATCTGCACCACTGGAAAGCTCCCTGTGCCATGGCCAACACTGGTCACTACCCTGACCTATGGCGTGCAGTGCTTTT
CCAGATACCCAGACCATATGAAGCAGCATGACTTTTTCAAGAGCGCCATGCCCGAGGGCTATGTGCAGGAGAGAACCATC
TTTTTCAAAGATGACGGGAACTACAAGACCCGCGCTGAAGTCAAGTTCGAAGGTGACACCCTGGTGAATAGAATCGAGCT
GAAGGGCATTGACTTTAAGGAGGATGGAAACATTCTCGGCCACAAGCTGGAATACAACTATAACTCCCACAATGTGTACA
TCATGGCCGACAAGCAAAAGAATGGCATCAAGGTCAACTTCAAGATCAGACACAACATTGAGGATGGATCCGTGCAGCTG
GCCGACCATTATCAACAGAACACTCCAATCGGCGACGGCCCTGTGCTCCTCCCAGACAACCATTACCTGTCCACCCAGTC
TGCCCTGTCTAAAGATCCCAACGAAAAGAGAGACCACATGGTCCTGCTGGAGTTTGTGACCGCTGCTGGGATCACACATG
GCATGGACGAGCTGTACAAGTGAGCGGCCGCAAGCTTGACTTCTTAACCCAACAGAAGGCTCGAGAAGGTATATTGCTGT
TGACAGTGAGCG**AAACAAGATGAAGAGCACCAAA**TAGTGAAGCCACAGATGTATTTGGTGCTCTTCATCTTGTTGTGCCT
ACTGCCTCGGACTTCAAGGGGCTAGAATTCGAACCGGTTTCCTAGGAGACATGATAAGATACATTGATGAGTTTGGACAA
ACCACAACTAGAATGCAGTGAAAAAAATGCTTTATTTGTGAAATTTGTGATGCTATTGCTTTATTTGTAACCATTATAAG
CTGCAATAAACAAGTTAACAACAACAATTGCATTCATTTTATGTTTCAGGTTCAGGGGGAGGTGTGGGAGGTTTTTTACG
CGTCAATTCTGTCATTTTACTAGGGTGATGAAATTCCCAAGCAACACCATCCTTTTCAGATAAGGGCACTGAGGCTGAGA
GAGGAGCTGAAACCTACCCGGCGTCACCACACACAGGTGGCAAGGCTGGGACCAGAAACCAGGACTGTTGACTGCAGCCC
GGTATTCATTCTTTCCATAGCCCACAGGGCTGTCAAAGACCCCAGGGCCTAGTCAGAGGCTCCTCCTTCCTGGAGAGTTC
CTGGCACAGAAGTTGAAGCTCAGCACAGCCCCCTAACCCCCAACTCTCTCTGCAAGGCCTCAGGGGTCAGAACACTGGTG
GAGCAGATCCTTTAGCCTCTGGATTTTAGGGCCATGGTAGAGGGGGTGTTGCCCTAAATTCCAGCCCTGGTCTCAGCCCA
ACACCCTCCAAGAAGAAATTAGAGGGGCCATGGCCAGGCTGTGCTAGCCGTTGCTTCTGAGCAGATTACAAGAAGGGACC
AAGACAAGGACTCCTTTGTGGAGGTCCTGGCTTAGGGAGTCAAGTGACGGCGGCTCAGCACTCACGTGGGCAGTGCCAGC
CTCTAAGAGTGGGCAGGGGCACTGGCCACAGAGTCCCAGGGAGTCCCACCAGCCTAGTCGCCAGACCGGGGATCCTCTAG
AGGATCCGGTACTCGAGGAACTGAAAAACCAGAAAGTTAACTGGTAAGTTTAGTCTTTTTGTCTTTTATTTCAGGTCCCG
GATCCGGTGGTGGTGCAAATCAAAGAACTGCTCCTCAGTGGATGTTGCCTTTACTTCTAGGCCTGTACGGAAGTGTTACT

TCTGCTCTAAAAGCTGCGGAATTGTACCCGCGGCCGCCACCATGGTGAGCAAGGGCGAGGAGGACAACATGGCCATCATC
AAGGAGTTCATGCGCTTCAAGGTGCACATGGAGGGCTCCGTGAACGGCCACGAGTTCGAGATCGAGGGCGAGGGCGAGGG
CCGCCCCTACGAGGGCACCCAGACCGCCAAGCTGAAGGTGACCAAGGGCGGCCCCCTGCCCTTCGCCTGGGACATCCTGT
CCCCTCAGTTCATGTACGGCTCCAAGGCCTACGTGAAGCACCCCGCCGACATCCCCGACTACTTGAAGCTGTCCTTCCCC
GAGGGCTTCAAGTGGGAGCGCGTGATGAACTTCGAGGACGGCGGCGTGGTGACCGTGACCCAGGACTCCTCCCTGCAGGA
CGGCGAGTTCATCTACAAGGTGAAGCTGCGCGGCACCAACTTCCCCTCCGACGGCCCCGTAATGCAGAAGAAGACCATGG
GCTGGGAGGCCTCCTCCGAGCGGATGTACCCCGAGGACGGCGCCCTGAAGGGCGAGATCAAGCAGAGGCTGAAGCTGAAG
GACGGCGGCCACTACGACGCCGAGGTCAAGACCACCTACAAGGCCAAGAAGCCCGTGCAGCTGCCCGGCGCCTACAACGT
CAACATCAAGCTGGACATCACCTCCCACAACGAGGACTACACCATCGTGGAACAGTACGAGCGCGCCGAGGGCCGCCACT
CCACCGGCGGCATGGACGAGCTGTACAAGTAAATGCATGACTTCTTAACCCAACAGAAGGCTCGAGAAGGTATATTGCTG
TTGACAGTGAGCG**AAACAAGATGAAGAGCACCAAA**TAGTGAAGCCACAGATGTATTTGGTGCTCTTCATCTTGTTGTGCC
TACTGCCTCGGACTTCAAGGGGCTAGAATTCGACTAGAGCTCGCTGATCAGCCTCGACTGTGCCTTCTAGTTGCCAGCCA
TCTGTTGTTTGCCCCTCCCCCGTGCCTTCCTTGACCCTGGAAGGTGCCACTCCCACTGTCCTTTCCTAATAAAATGAGGA
AATTGCATCGCATTGTCTGAGTAGGTGTCATTCTATTCTGGGGGGTGGGGTGGGGCAGGACAGCAAGGGGGAGGATTGGG
AAGACAATAGCAGGCATGCTGGGGAGAGATCTAGGAACCCCTAGTGATGGAGTTGGCCACTCCCTCTCTGCGCGCTCGCT
CGCTCACTGAGGCCGCCCGGGCAAAGCCCGGGCGTCGGGCGACCTTTGGTCGCCCGGCCTCAGTGAGCGAGCGAGCGCGC
AGAGAGGGAGTGGCCAACCCCCCCCCCCCCCCCCCTGCAGCCCTGCATTAATGAATCGGCCAACGCGCGGGGAGAGGCGG
TTTGCGTATTGGGCGCTCTTCCGCTTCCTCGCTCACTGACTCGCTGCGCTCGGTCGTTCGGCTGCGGCGAGCGGTATCAG
CTCACTCAAAGGCGGTAATACGGTTATCCACAGAATCAGGGGATAACGCAGGAAAGAACATGTGAGCAAAAGGCCAGCAA
AAGGCCAGGAACCGTAAAAAGGCCGCGTTGCTGGCGTTTTTCCATAGGCTCCGCCCCCCTGACGAGCATCACAAAAATCG
ACGCTCAAGTCAGAGGTGGCGAAACCCGACAGGACTATAAAGATACCAGGCGTTTCCCCCTGGAAGCTCCCTCGTGCGCT
CTCCTGTTCCGACCCTGCCGCTTACCGGATACCTGTCCGCCTTTCTCCCTTCGGGAAGCGTGGCGCTTTCTCAATGCTCA
CGCTGTAGGTATCTCAGTTCGGTGTAGGTCGTTCGCTCCAAGCTGGGCTGTGTGCACGAACCCCCCGTTCAGCCCGACCG
CTGCGCCTTATCCGGTAACTATCGTCTTGAGTCCAACCCGGTAAGACACGACTTATCGCCACTGGCAGCAGCCACTGGTA
ACAGGATTAGCAGAGCGAGGTATGTAGGCGGTGCTACAGAGTTCTTGAAGTGGTGGCCTAACTACGGCTACACTAGAAGG
ACAGTATTTGGTATCTGCGCTCTGCTGAAGCCAGTTACCTTCGGAAAAAGAGTTGGTAGCTCTTGATCCGGCAAACAAAC
CACCGCTGGTAGCGGTGGTTTTTTTGTTTGCAAGCAGCAGATTACGCGCAGAAAAAAAGGATCTCAAGAAGATCCTTTGA
TCTTTTCTACGGGGTCTGACGCTCAGTGGAACGAAAACTCACGTTAAGGGATTTTGGTCATGAGATTATCAAAAAGGATC
TTCACCTAGATCCTTTTAAATTAAAAATGAAGTTTTAAATCAATCTAAAGTATATATGAGTAAACTTGGTCTGACAGTTA
CCAATGCTTAATCAGTGAGGCACCTATCTCAGCGATCTGTCTATTTCGTTCATCCATAGTTGCCTGACTCCCCGTCGTGT
AGATAACTACGATACGGGAGGGCTTACCATCTGGCCCCAGTGCTGCAATGATACCGCGAGACCCACGCTCACCGGCTCCA
GATTTATCAGCAATAAACCAGCCAGCCGGAAGGGCCGAGCGCAGAAGTGGTCCTGCAACTTTATCCGCCTCCATCCAGTC
TATTAATTGTTGCCGGGAAGCTAGAGTAAGTAGTTCGCCAGTTAATAGTTTGCGCAACGTTGTTGCCATTGCTACAGGCA
TCGTGGTGTCACGCTCGTCGTTTGGTATGGCTTCATTCAGCTCCGGTTCCCAACGATCAAGGCGAGTTACATGATCCCCC
ATGTTGTGCAAAAAAGCGGTTAGCTCCTTCGGTCCTCCGATCGTTGTCAGAAGTAAGTTGGCCGCAGTGTTATCACTCAT
GGTTATGGCAGCACTGCATAATTCTCTTACTGTCATGCCATCCGTAAGATGCTTTTCTGTGACTGGTGAGTACTCAACCA
AGTCATTCTGAGAATAGTGTATGCGGCGACCGAGTTGCTCTTGCCCGGCGTCAATACGGGATAATACCGCGCCACATAGC
AGAACTTTAAAAGTGCTCATCATTGGAAAACGTTCTTCGGGGCGAAAACTCTCAAGGATCTTACCGCTGTTGAGATCCAG
TTCGATGTAACCCACTCGTGCACCCAACTGATCTTCAGCATCTTTTACTTTCACCAGCGTTTCTGGGTGAGCAAAAACAG
GAAGGCAAAATGCCGCAAAAAAGGGAATAAGGGCGACACGGAAATGTTGAATACTCATACTCTTCCTTTTTCAATATTAT
TGAAGCATTTATCAGGGTTATTGTCTCATGAGCGGATACATATTTGAATGTATTTAGAAAAATAAACAAATAGGGGTTCC
GCGCACATTTCCCCGAAAAGTGCCACCTGACGTCTAAGAAACCATTATTATCATGACATTAACCTATAAAAATAGGCGTA
TCACGAGGCCCTTTCGTCTCGCGCGTTTCGGTGATGACGGTGAAAACCTCTGACACATGCAGCTCCCGGAGACGGTCACA
GCTTGTCTGTAAGCGGATGCCGGGAGCAGACAAGCCCGTCAGGGCGCGTCAGCGGGTGTTGGCGGGTGTCGGGGCTGGCT
TAACTATGCGGCATCAGAGCAGATTGTACTGAGAGTGCACCATATGCGGTGTGAAATACCGCACAGATGCGTAAGGAGAA
AATACCGCATCAGGAAATTGTAAACGTTAATATTTTGTTAAAATTCGCGTTAAATTTTTGTTAAATCAGCTCATTTTTTA
ACCAATAGGCCGAAATCGGCAAAATCCCTTATAAATCAAAAGAATAGACCGAGATAGGGTTGAGTGTTGTTCCAGTTTGG
AACAAGAGTCCACTATTAAAGAACGTGGACTCCAACGTCAAAGGGCGAAAAACCGTCTATCAGGGCGATGGCCCACTACG
TGAACCATCACCCTAATCAAGTTTTTTGGGGTCGAGGTGCCGTAAAGCACTAAATCGGAACCCTAAAGGGAGCCCCCGAT
TTAGAGCTTGACGGGGAAAGCCGGCGAACGTGGCGAGAAAGGAAGGGAAGAAAGCGAAAGGAGCGGGCGCTAGGGCGCTG
GCAAGTGTAGCGGTCACGCTGCGCGTAACCACCACACCCGCCGCGCTTAATGCGCCGCTACAGGGCGCGTCGCGCCATTC
GCCATTCAGGCTACGCAACTGTTGGGAAGGGCGATCGGTGCGGGCCTCTTCGCTATTACGCCAGGCTGCAGGGGGGGGGG
GGGGGG

**shCtrl:** **AAACAAGATGAAGAGCACCAAA**

**Table S1 | Summarized donor information**

Young (N=15)

| ID | Age (y) | Gender | Post-mortem interval (time from death to tissue freezing) (h:min) |
|---|---|---|---|
| HE19 | 17 | M | 30:00 |
| HE51 | 22 | M | 24:00 |
| HE04 | 24 | M | 20:41 |
| HE52 | 28 | M | 16:20 |
| HE75 | 30 | F | 17:30 |
| HE25 | 31 | M | 29:00 |
| HE27 | 34 | F | 26:41 |
| HE67 | 35 | F | 49:45 |
| HE59 | 37 | M | 51:30 |
| HE35 | 39 | M | 24:00 |
| HE58 | 42 | M | 26:45 |
| HE53 | 42 | M | 26:00 |
| HE50 | 43 | M | 24:45 |
| HE55 | 45 | M | 28:41 |
| HE70 | 55 | F | 30:15 |

Old (N=14)

| ID | Age (y) | Gender | Post-mortem interval (time from death to tissue freezing) (h:min) |
|---|---|---|---|
| HE38 | 73 | F | 30:30 |
| HE16 | 74 | M | 34:35 |
| HE09 | 76 | M | 30:00 |
| HE07 | 77 | M | 24:10 |
| HE17 | 78 | M | 27:15 |
| HE13 | 80 | F | 15:00 |
| HE23 | 82 | M | 26:50 |
| HE21 | 83 | F | 20:45 |
| HE24 | 83 | F | 30:20 |
| HE29 | 87 | M | 21:50 |
| *HE08 | 90 | M | 31:40 |
| HE15 | 91 | F | 23:05 |
| HE37 | 93 | M | 21:20 |
| **HE74 | 94 | M | 21:25 |

* Cataract observed during eye enucleation

** Contralateral eye was diagnosed with AMD.

**Table S2 | Statistics for figure 2B**

| Comparisons | | | Adjusted p-values | | |
|---|---|---|---|---|---|
| | | | ***Hif1a*** | ***Hif2a*** | ***Slc2a1*** |
| shCtrl normoxia | vs. | shHif1 normoxia | <0.0001 | 0.9862 | 0.1316 |
| shCtrl normoxia | vs. | shHif2 normoxia | 0.9846 | <0.0001 | 0.9941 |
| shCtrl normoxia | vs. | shCtrl hypoxia | 0.9981 | 0.7104 | <0.0001 |
| shCtrl normoxia | vs. | shHif1 hypoxia | <0.0001 | <0.0001 | <0.0001 |
| shCtrl normoxia | vs. | shHif2 hypoxia | 0.7616 | <0.0001 | <0.0001 |
| shHif1 normoxia | vs. | shHif2 normoxia | <0.0001 | <0.0001 | 0.2869 |
| shHif1 normoxia | vs. | shCtrl hypoxia | <0.0001 | 0.3663 | <0.0001 |
| shHif1 normoxia | vs. | shHif1 hypoxia | >0.9999 | <0.0001 | <0.0001 |
| shHif1 normoxia | vs. | shHif2 hypoxia | <0.0001 | <0.0001 | <0.0001 |
| shHif2 normoxia | vs. | shCtrl hypoxia | 0.8892 | <0.0001 | <0.0001 |
| shHif2 normoxia | vs. | shHif1 hypoxia | <0.0001 | <0.0001 | <0.0001 |
| shHif2 normoxia | vs. | shHif2 hypoxia | 0.9800 | 0.9650 | <0.0001 |
| shCtrl hypoxia | vs. | shHif1 hypoxia | <0.0001 | <0.0001 | <0.0001 |
| shCtrl hypoxia | vs. | shHif2 hypoxia | 0.5313 | <0.0001 | 0.0013 |
| shHif1 hypoxia | vs. | shHif2 hypoxia | <0.0001 | <0.0001 | <0.0001 |
